# Beyond soil health: the trophic effects of cover crops shape predator communities

**DOI:** 10.1101/2020.03.28.013409

**Authors:** Carson Bowers, Michael D. Toews, Jason M. Schmidt

## Abstract

Maintaining habitat throughout the season in annual cropping systems provides resource stability for arthropod communities. Stabilizing resource availability should lead to diverse predatory communities and their associated ecosystem services such as biological control. There is a need for studies to test change in predator communities due to habitat provisioning and estimate associated food web responses. Here we quantified predator community structure and used molecular gut-content analysis to reconstruct predator food webs in response to winter cover crops (i.e. cereal and legume based) in a cotton agroecosystem. Predators were collected from experimental field plots during each major crop development stage in 2017 and 2018, and PCR was used to estimate predator roles and responses to cover crop treatments. Cotton planted into a rye cover crop residue promoted unique predator communities in the early and mid-season as compared to no-cover fields. Correspondingly, we observed dissimilar prey consumption among cover crop treatments. While predators consumed incidental pests at high frequencies (e.g. aphids), predation on key pests by natural enemies in this system was lacking. The use of winter cover crops and reduced tillage practices increased the consumption of alternative prey by natural enemies on seedling cotton, encouraging high predator diversity that aligns temporally with potential early season pest outbreaks. Therefore, cover crops should be further integrated into integrated pest management strategies.

## 1. Introduction

Annual cropping systems transition through cycles of vegetation growth and removal, affecting the prevalence and distribution of higher trophic level species (Wardle et al., 1999; Hanson et al., 2017). To dampen disturbance between cycles, maintaining habitat in annual systems provides spatial refuges that support prey and non-prey resources needed by predatory arthropods (Blüthgen et al., 2016). In particular, winter cover crops and increasing diversity of plant communities often results in a corresponding increase in the abundance and diversity of predators (Letourneau et al., 2011) and is thought to be correlated with the delivery of ecosystem services such as biological control (Gurr et al., 2017; Landis, 2017). However, elevated predator diversity may hinder service delivery by increasing negative interactions between beneficial species (Symondson et al., 2002) or through increasing the availability of preferred non-pest prey (Koss and Snyder 2005; Holt and Huxel, 2007). Consequently, simply improving diversity in agricultural systems is often not sufficient to reduce pest pressure and/or crop damage (Tscharntke et al., 2016).

The function of communities is often dependent on how environmental ‘filters’ such as habitat availability shape the composition of functional groups, and their corresponding interactions (Perovic et al., 2017). Past studies in agriculture focused on taxonomic differences to understanding diversity relationships with ecosystem functions (reviewed by Perovic et al. 2018), and studies suggest variable responses of species to habitat manipulation without noticeable changes in community function (Loreau & Mazancourt, 2013; Tilman et al., 1997), or species interactions (Ma et al., 2019). An evolved view argues for sorting species into functional groups based on combined traits (e.g. size, activity, habitat preferences, reproductive behavior, predation rates etc.) rather than the taxonomic identity of individual species (McEvoy 2018; Perovic et al., 2017; Gagic et al., 2015). Therefore, investigating the effectiveness of habitat manipulation for biological control service delivery requires quantifying changes in the structure of predator communities and associated interactions with economically significant prey (Gagic et al., 2015).

Estimating interactions among multiple species in a system, trophic interaction networks, are becoming more common as a proxy for predator function in agroecosystems (Poisot et al., 2013; Staudacher et al., 2018). Applying trophic networks to agroecosystems allows us to understand how predator-prey interactions are shaped by management practices and habitat dynamics of cropping systems (e.g. Tylianakis et al., 2007). Currently, some of the best empirical studies of habitat complexity influencing interactions (Finke & Denno, 2002; Hughes & Grabowski, 2006) are based on inference from changes in the abundances of a single or a few key prey and predators in field or cage studies (i.e. increased predator numbers and reduced pest numbers). However, generalist predators are common in many agroecosystems, and realized interaction frequencies are likely complex (Symondson et al., 2002; Furlong, 2015). Molecular gut-content analysis (MGCA) is an effective tool for identifying predation frequency under natural conditions (Eitzinger et al., 2018; Harwood et al., 2007; Szendrei et al., 2014; Ingrao et al., 2017; Roubinet et al., 2017). MGCA can provide estimates of interactions between entire predator communities, pests, and alternative prey in response to habitat manipulations and seasonal changes in crop growth.

Here we test the hypothesis that early season habitat provisioning and temporal dynamics of crop growth influence the composition of arthropod natural enemies and their interactions with prey in an annual cropping system. We use rye (*Secale cereale* L.) and crimson clover (*Tifolium incarnatum* L.) winter cover crops grown in a cotton agroecosystem to enhance early season habitat complexity compared to conventional cotton production. We build on the results of Bowers et al. (2020), where we found increased abundance and diversity of natural enemies in early season cover crop habitats. In the current study, we assess how changes in early season resource use may explain enhanced predator presence in response to cover cropping. We expected cover crop residue to provide both structural habitat (Holland et al., 2016) and fuel for detritus-based food webs (Chauvin et al., 2015) in the early season. Thus, habitat differences of cover crop residue and cotton at different development stages may result in a measurable change in community structure and resource use (i.e. consumption of prey). We predicted that 1) cotton grown into cover crops will support distinct natural enemy communities compared to without the use of a cover crop; 2) that predator communities will vary in their consumption of pest and non-pest prey when a cover crop is used with network metrics that indicate greater stability and more generalized feeding habitats; and 3) that community composition and trophic interactions will vary across the season (i.e. from early season cover residue to late season cotton). If cover crops shape predator communities and functional characteristics (i.e. predation), then building infield habitat may prove more effective in early season pest regulation than reliance on pesticides.

## 2. Materials and Methods

### 2.1. Experimental design and study site

We investigated the effects of cover crop habitat on predator community structure and function by establishing experimental plots in the fall of 2016 and 2017 at the UGA Southeast Georgia Research and Education Center at Midville, GA (Burke County, 32°52′15.6″N 82°13′12.0″W). Cover crop plots (0.4 ha) were established in a completely randomized block design and replicated 4 times for a total of 12 plots each year (Bowers et al., 2020). A conventionally managed control (no-cover) was maintained weed free throughout the off-season while crimson clover (27 kg/ha) and rye (67 kg/ha) cover crops were planted early November and chemically terminated and rolled 2 weeks before cotton planting. Cotton was planted into cover crop plots May 5, 2017 and April 28, 2018 using a Unverferth strip till rig, while no-cover plots were disked followed by a rip and bed pass. Plots were irrigated during cover crop growth and the cotton growing season, and no foliar insecticides were applied throughout the study (for full details please see Bowers et al., 2020).

### 2.2. Arthropod collection

Canopy and ground dwelling arthropods were sampled using a modified reverse-flow leaf blower within a 1m^2^ area quadrat (Bowers et al., 2020). In the previous study we presented the abundance and diversity responses. Here we analyzed the communities and trophic structure of predators counted in Bowers et al., 2020. On six dates the plots were sampled in three locations randomly selected (at minimum 10 m from any edge). All habitat (ground, cover residue, and cotton) within the 1 m^2^ quadrat was suctioned (~1 min/sample) until there was no visual arthropod activity detected (Bowers et al., 2020). All samples were placed in plastic bags and directly onto ice and within hours transferred to a −20°C freezer. Predators were then transferred to vials containing 96% ethanol and preserved at −20°C until identification and molecular analysis.

### 2.3. DNA Extraction

During sorting and identification, all intact, undamaged predators were prepared for molecular gut-content analysis. Predators were cleansed with 10% bleach solution, molecular grade H_2_O, and 95% ethanol, dried, and placed in sterile micro-centrifuge vials prior to DNA extraction to minimize environmental DNA contamination (Ingrao et al., 2017). Clean whole-body predators were pulverized for 30sec in 180 μl PBS using an agitator (Tissue Lyser II; QIAGEN, Chatsworth, CA, USA) and 3 mm stainless steel grinding balls (OPS Diagnostics), and total DNA was extracted and purified using QIAGEN DNeasy® 96 Blood and Tissue kit following manufacturer protocols for animal tissue extraction (QIAGEN, Chatsworth, CA, USA), with final elution of 75 μl AE buffer. Each 96-well plate included a negative control to test for DNA carry-over during the extraction process. Extracted predator DNA was stored at −20°C for later use in PCR.

### 2.4. Molecular gut-content analysis

Extracted predator DNA was screened for the presence of pest and alternative prey DNA using three separate PCR reactions, including two multiplex reactions using the Qiagen Multiplex PCR kit (Qiagen, Hilden, Germany) and a singleplex reaction; all utilizing established primer sets (Supporting Information Table S1). All reactions and primer mixtures were followed standard procedures for primer and PCR optimization (Staudacher et al., 2016, King et al., 2008; Sint et al., 2014). PCR reactions were designed to test for important economic pests (Lahiri et al., 2018; Naranjo 2018; Gowda et al., 2016), as well as occasional pests of cotton and common alternative prey taxa in agroecosystems (Nyffeler and Birkhofer 2017; Supporting Information Figure S1). “Multi1” was used to test for the DNA of primary cotton pests: Tarnished plant bugs (*Lygus spp.*) (Hagler and Blackmer, 2013), Southern green stink bug *(Nezara viridula*), whiteflies (*Bemisia tabaci*) (Itou et al., 2013) and Thrips (*Franklinellia spp.*) (Staudacher et al., 2016). A new primer set was designed and optimized for southern green stink bugs following recommended protocols for primer design in Chapman et al. (2013), and Staudacher et al. (2016), see Supporting Information S1. “Multi2” was used to test for the presence of an incidental cotton pest and common alternative prey that do not usually have economic significance to cotton production, but may influence interactions with primary pests. Multi2 tested for aphids (Family: Aphididae) (Staudacher et al., 2016), collembolans (Order: Collembola) (Staudacher et al., 2016), and dipterans (Order: Diptera) (Staudacher et al., 2016) DNA.

All PCR Reactions were run with a Bio-Rad C1000 Touch © Thermal Cycler (Bio-Rad, Hercules, California USA). Multi1 PCR reactions (12.5 μl) contained 6.25 μl 2x Qiagen multiplex master mix, 1.25 μl 10x primer mix, 0.5 μl 5x Q-solution, 0.3 μl BSA, 2.7 μl PCR grade H_2_0, and 1.5 μl extracted predator DNA. For Multi1, PCR protocol was 95°C for 15 min, followed by 34 cycles of 94°C for 30 s, 60°C for 45 s, 72°C for 1 min, and a final extension of 72°C for 5 min. Multi 2 reactions (12.5 μl) were comprised of 6.25 2x Qiagen multiplex master mix, 1.25 μl 10x primer mix, 1 μl 5x Q-solution, 0.3 μl BSA, 2.2 μl PCR grade H_2_0, and 1.5 μl extracted predator DNA. Multi2 PCR protocols were as follows: 95°C for 15 min, and 35 cycles of 94°C for 30 s, 63°C for 1 min 30 s, 72°C for 30 s, and a final extension of 72°C for 10 min. A singleplex PCR reaction was used to screen predators for an additional incidental pest, the two-spotted spider mite (*Tetranychus urticae*) (Krey et al., 2017). Aphids and spider mites are common pests of several cropping systems but do not consistently reach economically significant levels on cotton in the region (Rosenheim et al., 1997), and are therefore treated here as incidental pests. Screening for spider mites was done with a PCR reaction (12.5 μl) containing 6.25 μl (TopTaq, Qiagen, Hilden, Germany), 0.31 μl BSA, 0.625 μl of primers (forward and reverse), 3.69 μl PCR grade H_2_0, and 1 μl of extracted predator DNA. Spider mite reactions followed the protocol: 94°C for 3 min, and 44 cycles of 94°C for 30 s, 57°C for 30 s, 72°C for 1 min, with a final extension of 72°C for 1 min. All PCR products were visualized using a QIAxel Advanced System for DNA analysis, and incidents of predation were determined as positive prey DNA (>0.075 RFUs; Sint et al. 2014). Positive controls for each PCR reaction were included to verify reaction success and amplicon size for multiplex PCR. Predator-prey interactions were compared among cover cropping treatments and time periods.

### 2.5. Data analysis

The composition of predators and their interactions with cotton pests and alternative prey were analyzed using PERMANVOA models (fun: adonis2; package: vegan) using cover crop treatment and time period (early, mid, late) as fixed effects, and plot as a blocking factor (i.e., strata=plot) to account for repeated sampling of the same plot across each season. PERMANOVA models were based on 999 permutations and Bray-Curtis similarity measures (Legendre & Legendre, 1998). In the case of interactions between cover crop treatment and time, treatment effects at each time point (early, mid, late season) were assessed using separate PERMANOVA models for each year with treatment as a fixed effect for each time point. To investigate the effects of cover crops on interaction networks, commonly used network metrics (e.g. connectance, web asymmetry, links per species, weighted NODF, Shannon diversity, H2, niche overlap, functional complementarity; Dunne et al., 2002; Almeida-Neto et al. 2010; Tylianakis et al., 2007) were compared among treatments for each sample year using linear mixed effect models (LMM) using time period as a random effect. Significant differences in network metrics among treatments were assessed using pairwise comparisons and adjusted using Tukeys method (fun = lsmeans, package = lsmeans). All statistical analyses were performed in R v 3.3.2 (R Core Team, 2018). See publicly available data at dyrad.org, and also our further description of data handling in the supporting information S2.

## 3. Results

### 3.1. Predator community composition

A total of 2,675 predators were collected from the field across two sample years. Cover crops significantly influenced the composition of predators, which was dependent on the time period within each season (Figure 1). In 2017, predator communities were significantly influenced by cover crop treatment (PERMANOVA: F_2,207_=13.97, p=0.001), time period (PERMANOVA: F_2,207_=16.16, p=0.001), and an interaction between treatment and time period (PERMANOVA: F_4,207_=6.58, p=0.001). The interaction is explained by significant dissimilarity in predator community composition in early (PERMANOVA: F_2,9_=6.53, p=0.002; Figure 1a) and mid-season (PERMANOVA: F_2,9_=1.71, p=0.007; Figure 1b) in 2017. During late 2017 season, predator communities were not significantly dissimilar among cover crop treatments (PERMANOVA: F_2,9_=1.95, p=0.174). Similarly, in 2018, we observed dissimilar predator communities in cover crop treatments (PERMANOVA: F_2,219_=4.44, p=0.001), time periods (PERMANOVA: F_2,219_=29.97, p=0.001), and significant cover crop by time interaction (PERMANOVA: F_4,219_=2.89, p=0.001). This interaction can be explained by the significant dissimilarity of predator communities in the early (PERMANOVA: F_2,9_=16.02, p=0.002; Figure 1d) and mid (PERMANOVA: F_2,9_=4.39, p=0.005; Figure 1e) season but lack of differences in late season communities (PERMANOVA: F_2,9_=1.28, p=0.225; Figure 1f).

**Figure 1.**
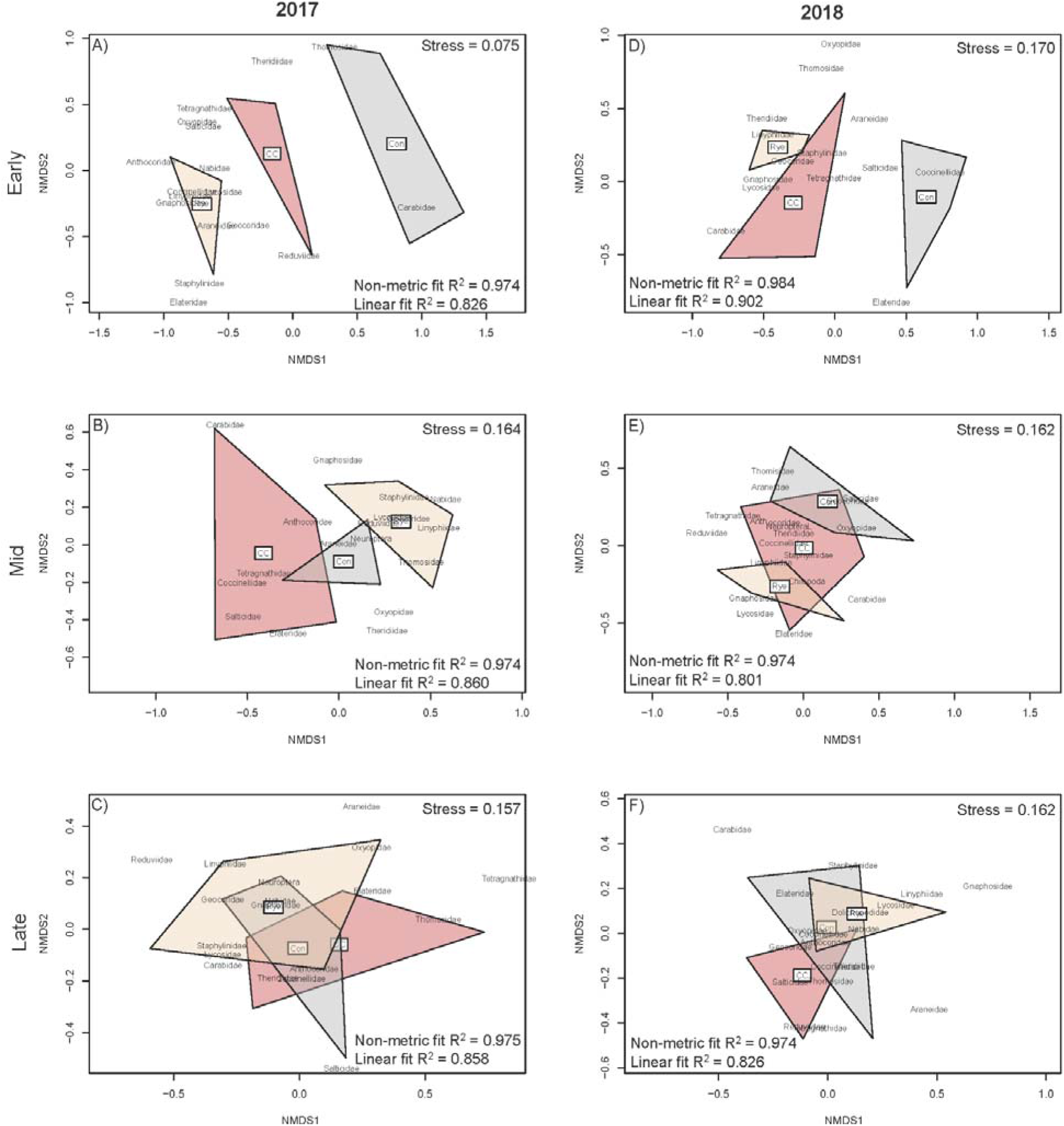
Non-metric multidimensional scaling (NMDS) for predator communities sampled from no cover (Con), Crimson clover (CC), and Rye (Rye) treatments in the early (Top), mid (mid), late (bottom) season in 2017 (left) and 2018 (right). Taxa included are present in more than 5% of total samples from each year.

In 2017, the use of crimson clover and rye cover crops resulted in an increased presence of several predator taxa including Geocoridae, Lycosidae, and Linyphiidae compared to conventionally managed cotton (Figure 2a) which harbored sparse predator communities comprised of mostly Carabidae beetles in the early season (Figure 2a; Supporting Information Figure S4). Ground dwelling spider taxa were common in mid-season rye, while conventional plots, and crimson clover appeared to attract more flying canopy predators such as Coccinellidae and *Orius* spp. (Family: Anthocoridae) (Supporting Information Figure S4). In 2018, The use of a cover crops resulted in a greater number of ground dwelling predator taxa including Lycosidae, Linyphiidae, and Gnaphosidae spiders, as well as Staphylinidae beetles compared to cotton grown without a cover crop, particularly when a rye cover crop was used (Figure 2b). This increase in ground dwelling predators in rye and crimson clover treatments continued into the mid-season, while conventional plots were shifting to dominance of canopy predators such as *Orius* and lacewings (Supporting Information Figure S4). In both years, the composition of late season predators was similar among treatments (Figure 1), though differing from early season communities through dominance of canopy predators such as *Orius* spp. and web building Theridiidae spiders depending on the sample year (Supporting Information Figure S4).

**Figure 2.**
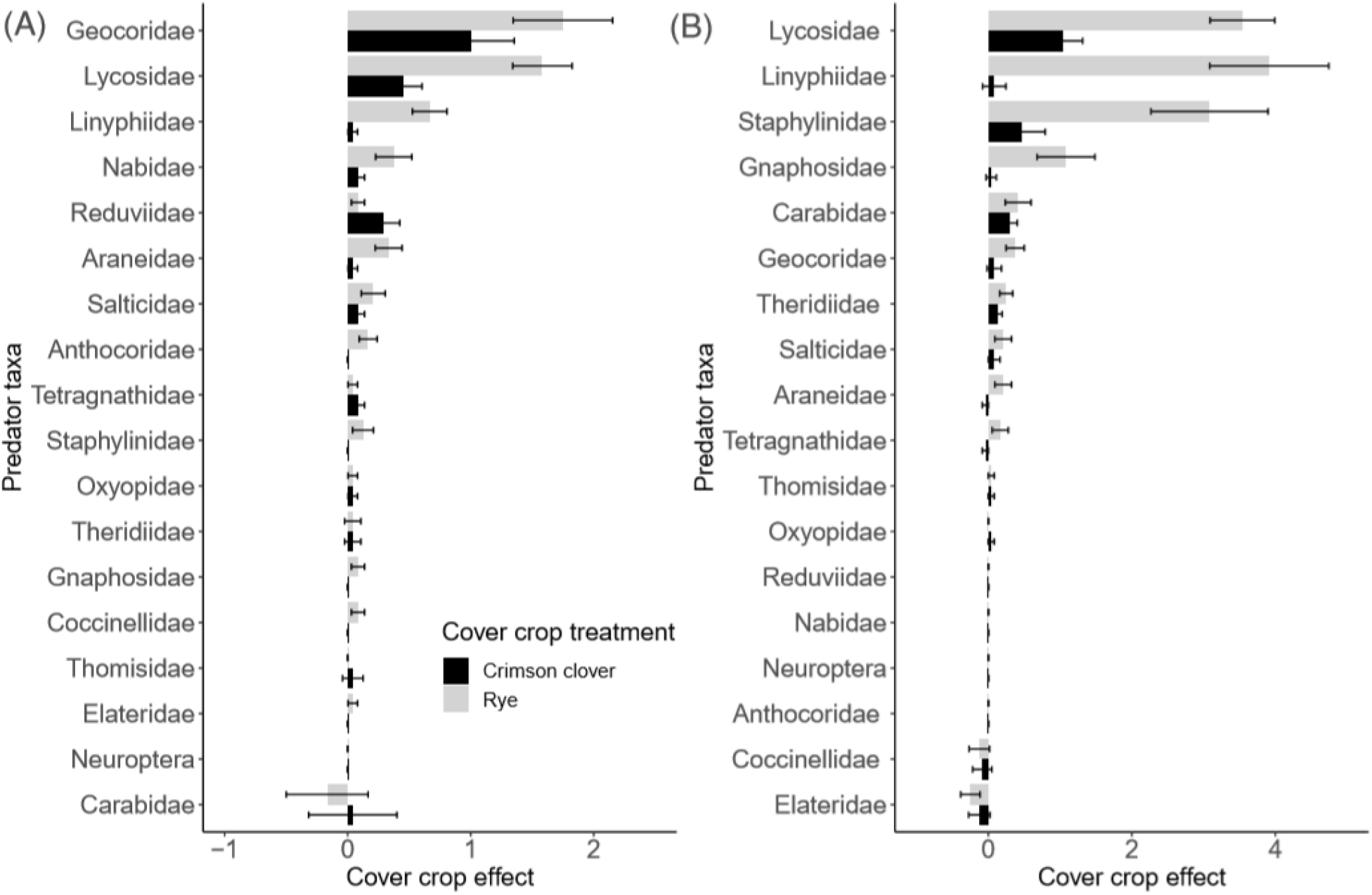
Mean difference in predator taxa abundance between cover crop treatments and no-cover conventional treatments in the early season (pre-emergence and seedling cotton) for 2017 and 2018. Error bars indicate standard error of the mean difference. Predator taxa are ranked by the strongest cover crop effect. See Supplemental for mid and late season cover crop effects and community composition by treatment and time period for each year.

### 3.2. Trophic interactions

A total of 2,454 predators were screened for the presence of pest and alternative prey DNA. The frequency of interactions between predators and prey in 2017 were significantly dissimilar by treatment (PERMANOVA: F_2,445_=7.96, p=0.001; Figure 3) and time period (PERMANOVA: F_2,445_=15.53, p=0.001; Figure 3). In 2018, we found that treatment (PERMANOVA: F_2,531_=14.11, p=0.001), time period (PERMANOVA: F_2,531_=13.30, p=0.001), as well a treatment by time period interaction (PERMANOVA: F_2,531_=3.35, p=0.003) significantly influenced trophic interactions. Consistently, our observed dissimilarity of interactions among treatments in both the early season (PERMANOVA: F_2,79_=6.09, p=0.005; Figure 3) and the mid-season (PERMANOVA: F_2,119_=3.09, p=0.009; Figure 3) but no difference in interactions in the late cotton growing season (PERMANOVA: F_2,175_=1.59, p=0.172; Figure 3). See supplemental for proportion positive of each prey by treatment and time period (Supporting Information Table S2, S3) and predator taxa (Supporting Information Table S4, S5).

**Figure 3.**
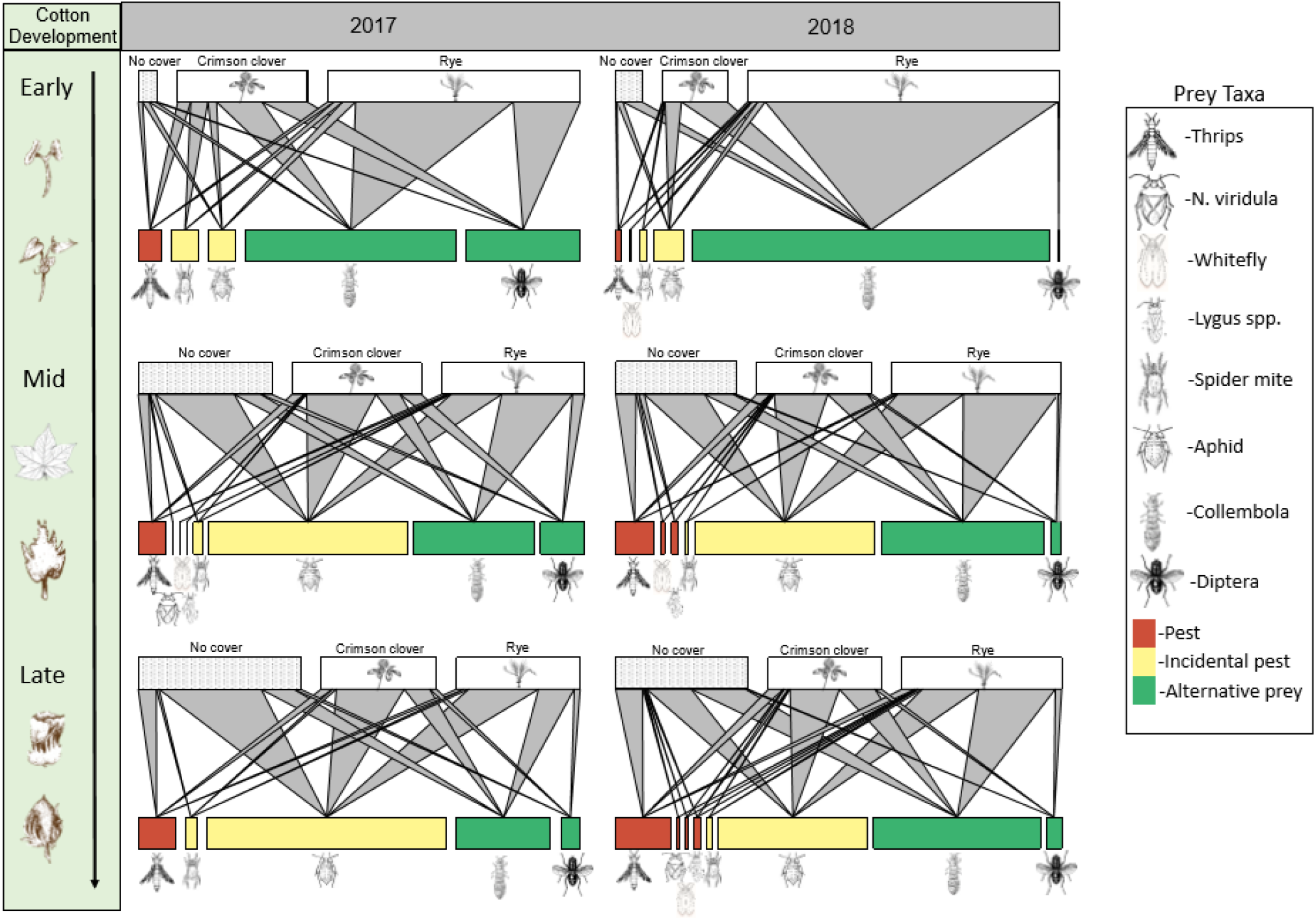
Food webs of predators pooled by cover crop treatment (No cover, Crimson clover, Rye) for early (top) mid (mid) and late (bottom) season in 2017 (left) and 2018 (right) with cover crop treatment represented by top bars with links to color coded prey bars on bottom of each web. The width of the bar indicating predator abundance relative to other treatments. Lower bars indicate prey taxa, with lines linking predators to prey and width of prey bars indicating relative frequency of positive PCR results by cover crop treatment. Prey taxa with no incidents of predation were excluded.

### 3.3. Network metrics

Cover crop use significantly influenced several network metrics in both 2017 and 2018 (Supporting Information Table S6, S7). In 2017, both weighted nestedness (NODF) and niche overlap (Supporting Information Table S6) were significantly influenced by cover crop treatment, with a higher degree of nestedness in rye compared to no cover and crimson clover treatments (Table 1), while crimson clover treatments had greater niche overlap compared to no cover treatments (Table 1). In 2018, only functional complementarity was influenced by cover crop treatment (Supporting Information Table S7), with rye treatments having significantly higher complementarity over either both crimson clover and no-cover treatments (Table 1).

**Table 1.**
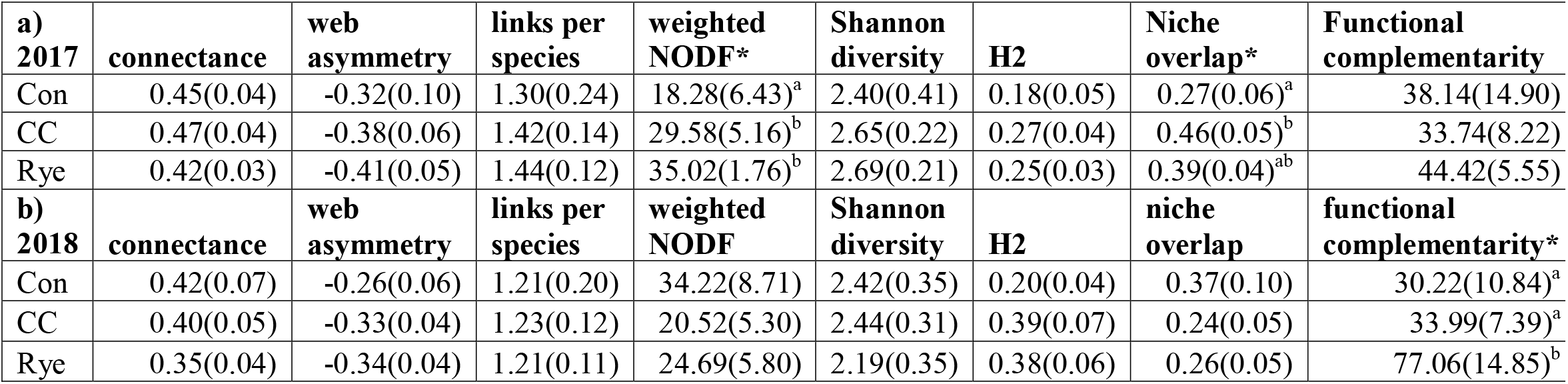
Summary of network metrics with the mean and SEM for each treatment in 2017 (a) and 2018 (b). Letters shown results of pairwise comparisons indicating significant differences (α=0.05) between treatments (Con=no cover; CC=Crimson clover).

## 4. Discussion

Cover crops facilitated high predator abundance and unique predator community structure in early season cotton. Trophic interaction networks responded to the addition of cover crop resources, and the lack of habitat in early season conventional cotton resulted in weak predator communities likely limited by prey availability. Thus, we demonstrate that among the many observed benefits (Hartwig and Ammon, 2002), cover crops fuel growth of early season arthropod communities through alternative prey consumption by predators.

Changes to the structure of predator communities in the early season was associated with habitat differences between cover crop treatments and conventional cotton production. Cotton grown into cover crop residue contained additional habitat structure and organic materials from plant biomass. In conventional, full tillage cotton, the only habitat available was seedling cotton. Cover crops selected for unique ground dwelling predator communities (Lycosidae and Staphylinidae; Figure 2), with rye particularly effective at harboring additional spider taxa such as Gnaphosidae and Linyphiidae (Figure 2). Ground dwelling insect predators and spiders respond positively to increasing habitat complexity (Mashavakure et al., 2019; Rivers et al., 2017) as a result of improved microhabitat and alternative prey availability. Without cover crop residue, seedling cotton lacks habitat and subsequently predators (Bowers et al., 2020), which likely increases the risk of early season pest damage (Cook et al., 2011; Toews et al., 2010). Therefore, cover crops improved the abundance and diversity of predatory taxa present (Bowers et al., 2020), and specifically amplified the growth of ground dwelling predator communities (Figure 2).

Using MGCA we assessed how feeding traits relevant to biological are shaped by the provisioning of early season cover cropping habitat. We identified increased consumption of alternative prey as a probable mechanism for altered structure and improved diversity of early season predator communities (Eveleigh et al., 2007). Predators collected from cotton grown into in cover crop residue were supported by alternative prey, indicated by increased frequency of detection of Collembola in the diets of predators across the season (Figure 3), and notably very high frequency in early season cotton grown into a rye cover crop. Additionally, nestedness, often interpreted as the asymmetry of interactions among specialist and generalist species (weighted NODF; Almeida-Neto et al., 2011), increased in cotton grown into cover crop residue. Thus, more rarely observed specialist species interacted with common species more often, increasing the connectedness of rare species to the entire network (Montoya et al. 2006). This may suggest the support of uncommon predator taxa by collembolan prey in these treatments, resulting in the observed increase in diversity in early season cover crop treatments. Conversely, predation by the small communities of predators in early season no-cover conventional treatments was biased toward aphid predation. Without cover crop residue present, predators are limited in their prey options on cotton seedlings, and likely extended their foraging range to the field margins or beyond in order meet their food needs (Rosenheim et al., 1997). This would explain low numbers of predators found within early season conventional treatments, while predators are maintained within the cropping area when cover crop residue is present and alternative prey resource availability is high. Therefore, cover crops altered the distribution and activity of predators by providing food resources within the crop area. In contrast to recent studies (Staudacher et al., 2018) we found no significant influence of cover crop treatment on the specialization (H2) of interaction networks. Where Stauadacher et al., 2018 found evidence for increasing generalization of interactions in more complex environments without changes to diversity, we found that changes in diversity associated with greater habitat complexity are not always accompanied by more generalized feeding networks. It is possible that the type of habitat and persistence of provisioned habitat influences the strength of change to either biodiversity or feeding interactions among predators and prey.

Despite the dynamic nature of annual cropping systems, few studies have investigated how the structure of predator communities and their interactions with prey change during seasonal crop growth. Trophic interaction networks respond to changes in the abundances of available species (Staniczenko et al., 2017), and seasonal changes in habitat. We provide clear early season evidence of dissimilar resource use between predators observed in cotton grown into a cover crop as compared to conventionally managed cotton. However, as the season progressed, habitat differences among treatments faded as the dominant vegetation shifts from cover crop residue to leafy cotton habitat, which resulted in comparable resource use by predator communities in the late season regardless of cover crop treatment (Figure 3). We also observed a shift in the structure of predator communities from early to late season from dominance of ground dwelling taxa in the early season to canopy predators (*Orius* spp., Theridiidae, Coccinellidae; Supporting Information Figure S3) in the late season. These shifts in composition are likely a result of the early provisioning of resources to ground dwelling taxa, and then the breakdown of these resources and subsequent increase in live cotton vegetation. The consumption of aphids increased among all treatments as the season progressed (Figure 3), likely indicating a shift in resource availability in the cropping area from detrital based resources provided by cover crop residue, to prey items which rely on live cotton vegetation. Recent studies which investigate how resource use changes across cropping seasons find similar increases in aphid predation later in the season (Roubinet et al., 2018). This indicates that changes in cropping habitat during the season results in changing predator function or feeding behavior at different periods of crop growth.

Although community level resource use was altered in the early season, realized contributions to biological control of key pests was not influenced by the use of winter cover crops. Indeed, we saw increased consumption of alternative prey by predator communities as well as greater degree of niche overlap and functional complementarity observed in crimson clover and rye respectively (Supporting Information Table S8 & S9). Our results suggest habitat driven changes to both the assembly of predatory communities and feeding behavior of predator communities toward detrital based food webs in the early season, and preference for “sugary” prey such as aphids later in the season. Therefore, behavioral and structural changes could not be linked to improved consumption of important economic pests, indicating a lack of predators with preferences toward key pests or ineffective current compositions of predators for pest suppression in both cover cropped or conventionally managed cotton. The only primary pest which predators tested positive in any substantial numbers were thrips. Unfortunately, the highest levels of thrips predation was seen in the mid and late season when thrips are of low risk to cotton (Toews et al., 2010; Mouden et al., 2017). Although the presence of alternative prey can improve the diversity of predators, our results of frequent feeding on alternative prey provides support for hindering the control of target pests (Koss and Snyder 2005; Holt and Huxel, 2007). The lack of predation on key pests across all treatments suggests vulnerabilities for the biological control of pests in this system.

Our interpretations of course have limitations. We are unable to assess how the availability of non-pest prey directly influenced the frequency of predation on key pests. Furthermore, the predation of pests such as thrips and whiteflies are likely conservative with current molecular methods, as the DNA of small prey such as thrips and whiteflies can degrade quickly within predator guts, reducing detectability to a few hours after consumption (Gomez-Polo et al., 2016). Although DNA decay trials may help adjust frequencies of predation, our study was not designed to precisely estimate the rates of DNA decay in such diverse communities (Greenstone et al. 2014). However, our results provide strong support for the hypothesis that cover crops contribute to pest management in cotton, and suggest that early season conventionally managed cotton is vulnerable to thrips outbreaks due to a lack of predator presence. We provide a comprehensive look at cotton predator communities and provide evidence for close associations between arthropod community structure, available habitat, and predator function. We show that the use of winter cover crops can reduce the risk of early pest outbreaks by encouraging the presence of stable predator communities through increased alternative prey consumption. Future work is needed to assess how altering the availability of prey influences prey preference of predators throughout the season and their control of key pests, by assessing prey population levels and quantifying antagonistic interactions between predators.

## Acknowledgements

Special thanks to Anthony Black for plot preparation and management. Additionally, the authors express thanks to Melissa Thompson, Zachary Wainwright, Abigail Borem, Emmalee Milner, Rachel Perez, Shereen Xavier for help with field collections and sample processing. Thanks to Megan McCoy for icon and food web figure artistic design. This work was supported, in part, by the University of Georgia, USDA-NIFA Multistate Hatch Project GEO00884-S1073, Georgia Cotton Commission project 15-156GA, and Cotton Incorporated. Mention of trade names or commercial products in this publication is solely for the purpose of providing specific information and does not imply recommendation or endorsement by the University of Georgia.

## Authors contributions

M.T. and J.S. obtained funding and conceived/designed the study together. C.B., M.T. and J.S. conducted field work/collected samples (with support of field assistants), C.B. identified specimens (morphologically), and performed laboratory work (with support of laboratory assistants). C.B. and J.S. analyzed the data/compiled tables and figures. C.B. and J.S. wrote the first draft of the manuscript, and M.T. contributed to editing and finalizing the submitted manuscript.

