## Supporting Information and Figures for "Beyond soil health: the trophic effects of cover crops shape predator communities"

**Table S1. Primer information**

| **Target** | **Name** | **Gene** | **5’-3’ sequence** | **Amplicon length (in multiplex)** | **PCR assay (conc., uM)** | **Primer source** |
| --- | --- | --- | --- | --- | --- | --- |
| *Lygus spp.* | AY25-lygusF  AY25-lygusR | COI | AGGATTTGGACTAATCTCAC  ATTACTCCAGTAAGACCTCCT | 324 | Multi 1 (0.2) | Hagler and Blackmer, 2013 |
| *Bemisia tabaci* | Bt-F  Bt-R | COI | TTGGTGCTCCTGACATAGCTT  TAAGCCTCTATGAGTTAATCTTAAA | 165 | Multi 1 (0.5) | Itou et al., 2013 |
| *Frankliniella spp.* | S477-thrips  A481-thrips | 18S | CGGTGTCAAACTGACGCGA  GCCCCCGCCTGTCTCC | 284 | Multi 1 (0.2 F, 0.4 R) | Staudacher et al., 2016 |
| *Nezara viridula* | SG-223F  SG-223R | COI | GCAGAATCTGGAGCAGGAACA  CTGTGATCCCAACTGAGCAAAC | 219 | Multi 1 (0.2) | Supplemental 1 |
| Aphids | S423-aphid1  A424-aphid2 | 18S | TGGTTCCTTAGATCGTACCCAAG  GCCGCGACGGGCC | 148 | Multi 2 (0.4) | Staudacher et al., 2016 |
| Collembola | S411-springtails1  A415-springtails1 | 18S | GCTCGTAGTTGGATYTCGGTTT  GAATTTCACCTCTAACGTCGCAG | 289 | Multi 2 (0.1) | Staudacher et al., 2016 |
| Diptera | S414-dipterans1  A416-dipterans2 | 18S | CCTATCAACTATTGATGGTAGTRTCKWGGA  GAAGCACAARWTCAACTWCGAACG | 341 | Multi 2 (0.5) | Staudacher et al., 2016 |
| Spider mite | Turtic-181-F  Turtic-396-R | COI | AGGATTTGGAAATTGATTGA  AAAATTATTATTTCAATAGAGGAAGAC | ~215 | Spider mite (0.6) | Krey et al., 2017 |

Figure S1. Gel images for multiplex PCR results. Shown is results for Multi 1 (left) and Multi 2 (right) including single bands for each prey taxa and positive controls with bands for all prey taxa screened. Bands shown include whitefly (165 bp), southern green stink bug (219 bp), thrips (284 bp) and lygus (324 bp) for multi 1. Multi 2 results shown include aphid (149 bp), Collembola (289 bp), Diptera (341 bp) as well as additional bands for burrow bug (254 bp) and lacewing (390 bp) although they were not of interest for this study.

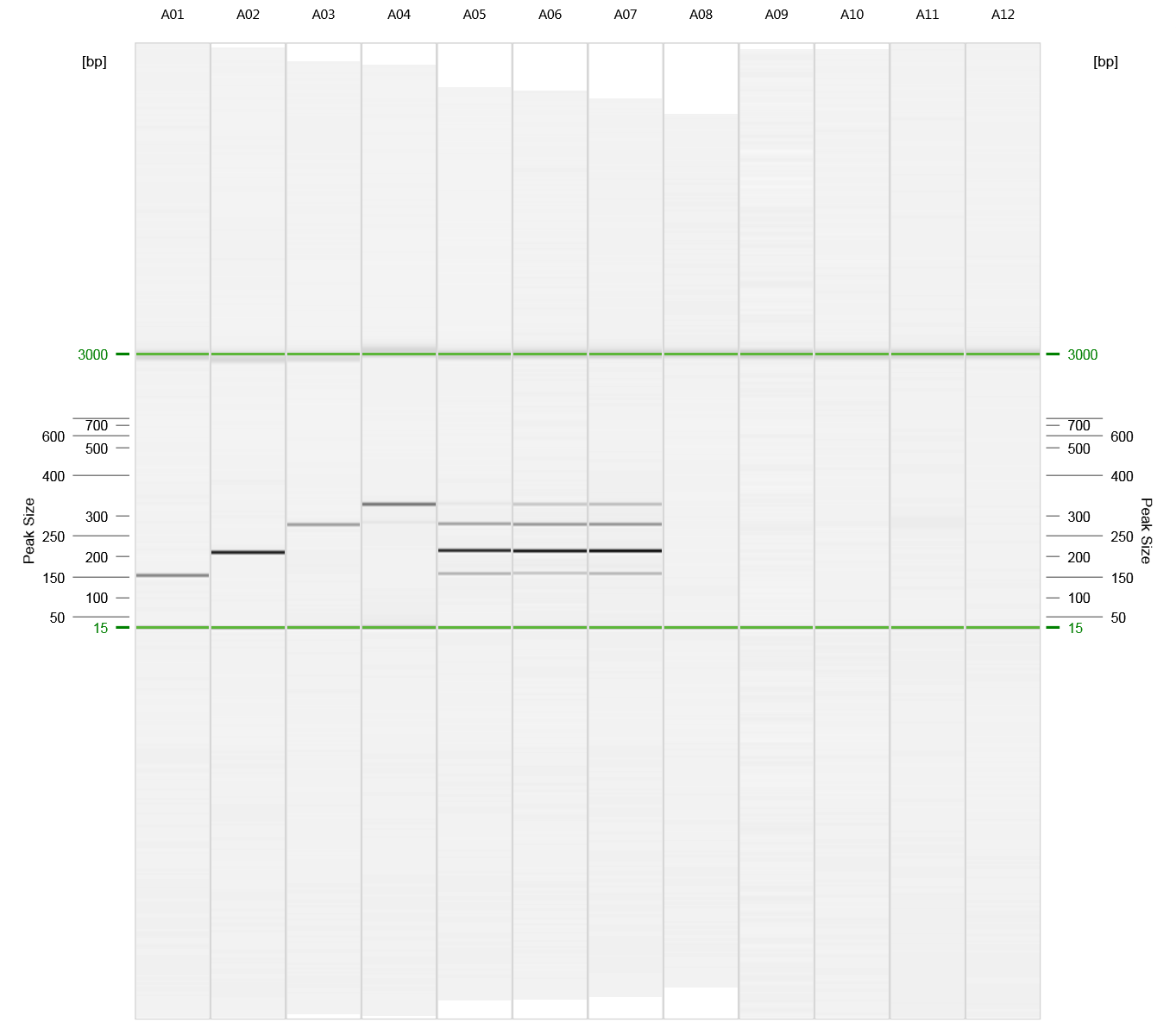

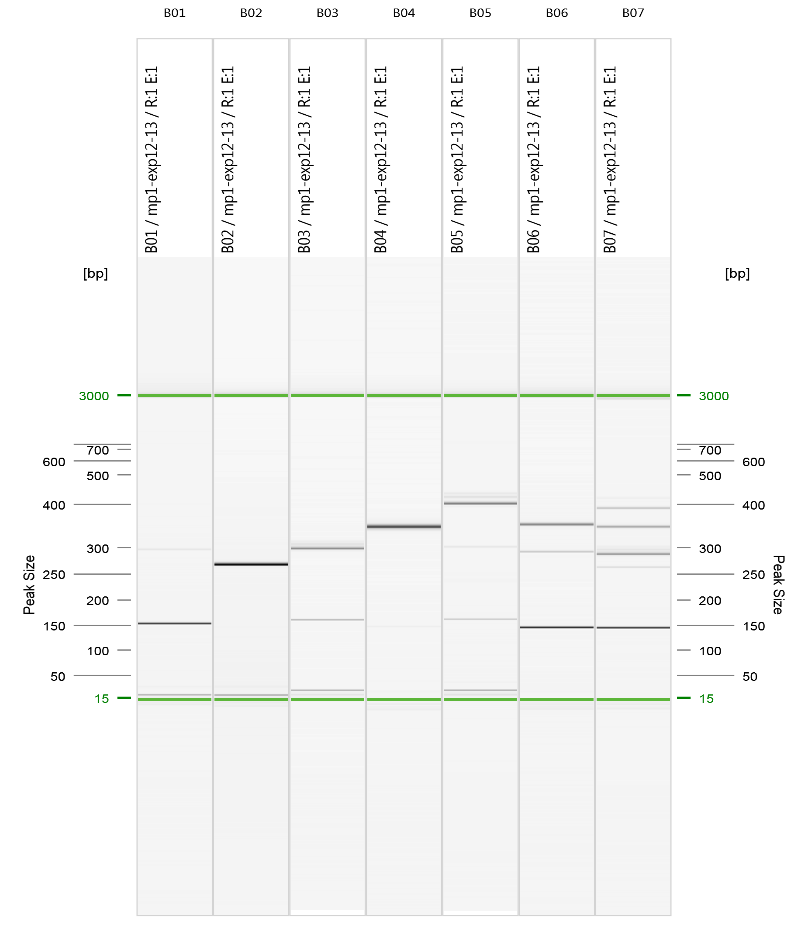

*S1. Primer Design* – Molecular gut-content analysis (MGCA) primer design for stink bugs and PCR conditions and modifications for other assays used from the literature.

We optimized a set of COI based primers for screening for *N.viridula* following best practices of creating new multiplex PCRs for diagnostic PCR in gut content analyses (King et al. 2008, 2011, Staudacher et al. 2016).  Using PRIMER 3 and PRIMER Blast on NCBI and GenBank sequences: KR037758.1 and KR044112.1, we selected a set of primers that appeared to not amplify other species in the GENBANK database, and provided an amplicon size compatible with molecular gut content analysis ~223bp:

SG-223F – 5’- GCAGAATCTGGAGCAGGAACA-3’

SG-223R – 5’- CTGTGATCCCAACTGAGCAAAC-3’

These were then tested in the laboratory for thermocycling conditions and cross-reactivity. The primers did not show cross-reactivity when screened against leg extractions of families we expected in our samples: 1) Araneae families: Araneidae, Corinnidae, Linyphiidae, Lycosidae, Salticidae; 2) Coleoptera families: Carabidae, Coccinellidae;

3) Diptera families: Dolichopodidae, Muscidae; 4) Hemiptera families: Miridae, Anthicoridae, Geocoridae.

The optimal thermocycling conditions were 15 min at 95°C, 35 cycles of 30s at 94°C, 90s at 60°C, 60s at 72°C, and 5 min at 72°C. We followed the same multiplex mix conditions using the Multiplex PCR Master Mix kit (Qiagen), and final primer concentrations were 2mM per primer in the final PCR mix, these were mixed prior to combining with PCR mix and added to form an overall primer concentration in the mix of 10mM as suggested by manufacture (Qiagen).

PCR conditions for other reactions used for Qiagen multiplex kit. For positive controls, we always used a mixed DNA solution containing standardized extraction concentrations of the targeted templates. It was possible to multiplex the Thrips (Staudacher et al. 2016), Lygus (Hagler et al. 2013), stink bug primers, and white fly primers (Itou et al. 2013) into a single reaction using the PCR conditions for Qiagen Multiplex Kit: 15 min at 95°C, 34 cycles of 94°C for 30 s, 60°C for 45 s, 72°C for 1 min, and a final extension of 72°C for 5 min, which formed four clear and consistent bands: 323bp for *Lygus*, 284 bp for thrips, 223bp for *N.viridula*, and 139bp for white flies (Supplementary Fig.1). For aphids, Diptera and Collembola (Multi 2; Staudacher et al. 2016) we followed the published protocols using same reagents and PCR protocols were as follows: 95°C for 15 min, and 35 cycles of 94°C for 30 s, 63°C for 1 min 30 s, 72°C for 30 s, and a final extension of 72°C for 10 min. For spider mites (Krey et al. 2017), we followed the same published protocol with Qiagen reagents.

*S2. Data management*

In the previous study, Bowers et al. 2020, we presented the overall counts of natural enemies, diversity of natural enemies and a full complement of data on production. Here we reconfigured the natural enemy data to exam community and functional attributes (i.e. feeding interactions with common pests, incidental pests, and alternative prey). Below we detail how we configured our datasets to target community analysis, and we also make these datasets available to whomever for future exploration.

Due to inter-annual variation in predator community composition and trophic interactions among treatments, each season (i.e. 2017, 2018) was analyzed separately. Predator taxa were pooled at the family level when investigating both community analysis and trophic interactions. To improve model adequacy, rare predators observed in less than five percent of samples across a single season were excluded from community analyses (following McCune & Grace 2002). To investigate the prey consumption of treatment specific arthropod communities included in the community composition analysis, only predators included in the community analysis were included in interaction matrices for each treatment or time period. For comparison of interaction network characteristics by treatment, network metrics generated (fun: networklevel; package: bipartite) were pooled by plot and sample date to maintain replication of treatments. To investigate the temporal changes in community composition and predator function in response to changes in habitat, predator communities were pooled by time period (early, mid, late) for each year, with each time period including two major cotton development stages (i.e. dates).

**Table S2. Predation by treatment and time period in 2017. Proportion of total predators (n) in each treatment that tested positive for each prey item at each time period (early, mid, late) is shown.**

| **trt** | **time** | **n** | **Thrips** | **Stink bug** | **whitefly** | **Lygus** | **Spider mite** | **Aphid** | **Collembola** | **Diptera** |
| --- | --- | --- | --- | --- | --- | --- | --- | --- | --- | --- |
| No cover | *Early* | 21 | 0.095 | 0.000 | 0.000 | 0.000 | 0.000 | 0.143 | 0.143 | 0.048 |
|  | *Mid* | 82 | 0.098 | 0.012 | 0.000 | 0.000 | 0.049 | 0.720 | 0.244 | 0.098 |
|  | *Late* | 213 | 0.127 | 0.000 | 0.000 | 0.000 | 0.019 | 0.714 | 0.230 | 0.047 |
| Crimson clover | *Early* | 77 | 0.078 | 0.000 | 0.000 | 0.000 | 0.104 | 0.052 | 0.273 | 0.273 |
|  | *Mid* | 82 | 0.110 | 0.000 | 0.000 | 0.012 | 0.012 | 0.634 | 0.220 | 0.195 |
|  | *Late* | 148 | 0.115 | 0.000 | 0.000 | 0.000 | 0.041 | 0.682 | 0.257 | 0.068 |
| Rye | *Early* | 155 | 0.019 | 0.000 | 0.000 | 0.000 | 0.026 | 0.039 | 0.477 | 0.206 |
|  | *Mid* | 114 | 0.035 | 0.000 | 0.009 | 0.000 | 0.018 | 0.333 | 0.482 | 0.079 |
|  | *Late* | 141 | 0.078 | 0.000 | 0.000 | 0.000 | 0.050 | 0.738 | 0.383 | 0.064 |

**Table S3. Predation by treatment and time period in 2018. Proportion of total predators (n) in each treatment that tested positive for each prey item at each time period (early, mid, late) is shown.**

| **trt** | **time** | **n** | **Thrips** | **Stink bug** | **whitefly** | **Lygus** | **Spider mite** | **Aphid** | **Collembola** | **Diptera** |
| --- | --- | --- | --- | --- | --- | --- | --- | --- | --- | --- |
| No cover | *Early* | 30 | 0.033 | 0.000 | 0.000 | 0.000 | 0.000 | 0.133 | 0.433 | 0.000 |
|  | *Mid* | 92 | 0.163 | 0.000 | 0.022 | 0.043 | 0.000 | 0.478 | 0.228 | 0.022 |
|  | *Late* | 183 | 0.180 | 0.011 | 0.005 | 0.011 | 0.000 | 0.295 | 0.361 | 0.016 |
| Crimson clover | *Early* | 75 | 0.013 | 0.000 | 0.000 | 0.000 | 0.013 | 0.147 | 0.413 | 0.000 |
|  | *Mid* | 81 | 0.123 | 0.000 | 0.012 | 0.012 | 0.012 | 0.481 | 0.358 | 0.037 |
|  | *Late* | 146 | 0.123 | 0.000 | 0.007 | 0.034 | 0.014 | 0.411 | 0.295 | 0.055 |
| Rye | *Early* | 332 | 0.006 | 0.000 | 0.003 | 0.000 | 0.012 | 0.015 | 0.596 | 0.003 |
|  | *Mid* | 125 | 0.024 | 0.000 | 0.000 | 0.000 | 0.008 | 0.384 | 0.552 | 0.016 |
|  | *Late* | 190 | 0.084 | 0.011 | 0.005 | 0.005 | 0.026 | 0.358 | 0.489 | 0.042 |

**Table S4. 2017 Proportion positive of each prey item by predator taxa (family level). Predators can test positive for more than 1 prey taxa, therefore the proportion of prey combined does not necessarily total 1. N is the total number of individuals screened from each predator taxa.**

| **Predator taxa** | **n** | **Thrips** | **Stink bug** | **whitefly** | **Lygus** | **Spider mite** | **Aphid** | **Collembola** | **Diptera** |
| --- | --- | --- | --- | --- | --- | --- | --- | --- | --- |
| Anthocoridae | 230 | 0.104 | - | - | - | 0.009 | 0.557 | 0.083 | 0.004 |
| Lycosidae | 111 | - | - | - | - | 0.090 | 0.441 | 0.874 | 0.036 |
| Coccinellidae | 95 | 0.084 | - | - | - | 0.011 | 0.958 | 0.242 | 0.074 |
| Geocoridae | 94 | 0.064 | - | - | - | 0.106 | 0.234 | 0.479 | 0.457 |
| Carabidae | 77 | - | - | - | - | 0.013 | 0.117 | 0.130 | 0.299 |
| Nabidae | 47 | 0.021 | - | - | - | - | 0.702 | 0.149 | 0.021 |
| Theridiidae | 46 | 0.043 | - | - | - | 0.065 | 0.696 | 0.304 | 0.022 |
| Araneidae | 39 | 0.128 | - | - | - | 0.026 | 0.462 | 0.333 | 0.154 |
| Neuroptera | 38 | 0.289 | - | - | - | 0.105 | 1.000 | 0.184 | 0.026 |
| Linyphiidae | 34 | 0.029 | - | - | - | - | 0.118 | 0.588 | - |
| Elateridae | 31 | 0.032 | - | - | - | - | 0.677 | 0.129 | 0.032 |
| Gnaphosidae | 30 | - | - | 0.033 | - | - | 0.133 | 0.667 | 0.033 |
| Salticidae | 28 | 0.393 | - | - | - | 0.036 | 0.464 | 0.429 | 0.036 |
| Reduviidae | 26 | 0.038 | 0.038 | - | - | - | 0.423 | 0.308 | 0.269 |
| Staphylinidae | 26 | 0.115 | - | - | - | 0.038 | 0.308 | 0.115 | - |
| Oxyopidae | 23 | 0.043 | - | - | 0.043 | - | 0.739 | 0.087 | 0.348 |
| Thomisidae | 17 | 0.412 | - | - | - | - | 0.353 | 0.412 | 0.176 |
| Tetragnathidae | 14 | 0.071 | - | - | - | - | 0.071 | 0.786 | 0.214 |
| Araneae immature | 12 | - | - | - | - | - | - | 0.333 | 0.083 |
| Dermaptera | 9 | 0.222 | - | - | - | 0.111 | 0.889 | 0.444 | 0.222 |
| Anthicidae | 1 | - | - | - | - | 1.000 | 1.000 | - | - |
| Clubionidae | 1 | 1.000 | - | - | - | - | 1.000 | - | - |
| Corinnidae | 1 | - | - | - | - | - | 1.000 | 1.000 | - |
| Dictynidae | 1 | - | - | - | - | - | 1.000 | 1.000 | - |
| Pisauridae | 1 | - | - | - | - | - | 1.000 | - | 1.000 |
| Tettigoniidae | 1 | 1.000 | - | - | - | - | 1.000 | - | 1.000 |

**Table S5. Proportion positive of each prey item by predator taxa (family level) in 2018. Predators can test positive for more than 1 prey taxa, therefore P prey combined does not necessarily total 1. N is the total number of individuals screened from each predator taxa.**

|  | **N** | **Thrips** | **Stink bug** | **whitefly** | **Lygus** | **Spider mite** | **Aphid** | **Collembola** | **Diptera** |
| --- | --- | --- | --- | --- | --- | --- | --- | --- | --- |
| Lycosidae | 217 | 0.009 | - | - | 0.014 | 0.023 | 0.157 | 0.949 | - |
| Theridiidae | 140 | 0.057 | 0.014 | - | 0.007 | 0.014 | 0.371 | 0.521 | - |
| Linyphiidae | 123 | 0.008 | - | - | - | 0.016 | 0.065 | 0.813 | - |
| Staphylinidae | 115 | - | - | - | - | - | 0.035 | 0.096 | 0.009 |
| Coccinellidae | 113 | 0.106 | - | - | - | - | 0.735 | 0.345 | - |
| Anthocoridae | 75 | 0.133 | - | - | - | - | 0.347 | 0.147 | - |
| Salticidae | 67 | 0.328 | 0.015 | 0.104 | 0.090 | 0.060 | 0.552 | 0.478 | - |
| Geocoridae | 43 | 0.116 | - | - | - | - | 0.093 | 0.140 | 0.023 |
| Thomisidae | 43 | 0.372 | 0.023 | - | 0.047 | - | 0.256 | 0.419 | - |
| Gnaphosidae | 41 | - | - | - | - | - | - | 0.463 | - |
| Elateridae | 35 | - | - | - | - | - | 0.029 | 0.143 | - |
| Chilopoda | 32 | - | - | - | - | - | 0.031 | 0.156 | - |
| Hemerobiidae | 29 | 0.241 | - | - | - | - | 0.862 | 0.207 | 0.069 |
| Carabidae | 23 | - | - | - | - | - | 0.130 | 0.043 | - |
| Araneidae | 22 | 0.091 | - | - | - | 0.045 | 0.182 | 0.318 | - |
| Oxyopidae | 22 | 0.227 | - | - | 0.045 | - | 0.500 | 0.091 | 0.091 |
| Dolichopodidae | 20 | 0.150 | - | - | - | - | 0.200 | 0.100 | 1.000 |
| Nabidae | 20 | - | - | - | - | - | 0.200 | 0.050 | - |
| Chrysopidae | 20 | 0.150 | - | - | - | - | 0.500 | 0.100 | - |
| Anthicidae | 14 | - | - | - | - | - | 0.071 | - | - |
| Tetragnathidae | 12 | - | - | - | - | - | 0.083 | 0.750 | - |
| Reduviidae | 8 | - | - | - | - | - | 0.250 | - | - |
| Dermaptera | 6 | - | - | - | - | - | - | 0.167 | - |
| Miturgidae | 5 | 0.200 | - | - | - | - | 0.600 | 0.400 | 0.200 |
| Anyphaenidae | 2 | - | - | - | - | - | 0.500 | 0.500 | - |
| Mimetidae | 2 | 0.500 | - | - | - | - | 0.500 | - | - |
| Pisauridae | 2 | 0.500 | - | - | - | - | 0.500 | 1.000 | - |
| Dictynidae | 1 | - | - | - | - | - | - | 1.000 | - |
| Philodromidae | 1 | - | - | - | - | - | 1.000 | 1.000 | - |
| Araneae Immature | 1 | - | - | - | - | - | - | - | - |

**Table S6. Results of linear mixed models of 2017 for network metrics in relation to cover crop treatment. F and P values for each metric are shown, with significant differences (α=0.05) shown in bold.**

|  | treatment |  |
| --- | --- | --- |
| Network Metric | F | P |
| Connectance | 0.547 | 0.592 |
| Web asymmetry | 1.475 | 0.265 |
| Links per species | 0.545 | 0.593 |
| Weighted NODF | 4.395 | **0.035** |
| Shannon diversity | 0.865 | 0.444 |
| H2 | 1.264 | 0.317 |
| Niche overlap | 3.501 | **0.056** |
| Functional complimentarity | 0.354 | 0.701 |

**Table S7. Results of linear mixed models of 2018 for network metrics in relation to cover crop treatment F and P values for each metric are shown, with significant differences (α=0.05) shown in bold.**

|  | treatment |  |
| --- | --- | --- |
| Network Metric | F | P |
| Connectance | 0.682 | 0.523 |
| Web asymmetry | 0.770 | 0.481 |
| Links per species | 0.0178 | 0.982 |
| Weighted NODF | 1.382 | 0.286 |
| Shannon diversity | 0.425 | 0.662 |
| H2 | 1.264 | 0.317 |
| Niche overlap | 0.919 | 0.420 |
| Functional complimentarity | 5.178 | **0.020** |

**Table S8. Summary of network metrics with the mean and SEM for each treatment in 2017. Letters shown results of pairwise comparisons indicating significant differences (α=0.05) between treatments or time periods.**

| **a)** | **connectance** | **web asymmetry** | **links per species** | **weighted NODF*** | **Shannon diversity** | **H2** | **Niche overlap*** | **Functional complementarity** |
| --- | --- | --- | --- | --- | --- | --- | --- | --- |
| Con | 0.45(0.04) | -0.32(0.10) | 1.30(0.24) | 18.28(6.43)^a^ | 2.40(0.41) | 0.18(0.05) | 0.27(0.06)^a^ | 38.14(14.90) |
| CC | 0.47(0.04) | -0.38(0.06) | 1.42(0.14) | 29.58(5.16)^b^ | 2.65(0.22) | 0.27(0.04) | 0.46(0.05)^b^ | 33.74(8.22) |
| Rye | 0.42(0.03) | -0.41(0.05) | 1.44(0.12) | 35.02(1.76)^b^ | 2.69(0.21) | 0.25(0.03) | 0.39(0.04)^ab^ | 44.42(5.55) |

**Table S9. Summary of network metrics with the mean and SEM for each treatment in 2018. Letters shown results of pairwise comparisons indicating significant differences (α=0.05) between treatments or time periods.**

| **b)** | **connectance** | **web asymmetry** | **links per species** | **weighted NODF** | **Shannon diversity** | **H2** | **niche overlap** | **functional complementarity*** |
| --- | --- | --- | --- | --- | --- | --- | --- | --- |
| Con | 0.42(0.07) | -0.26(0.06) | 1.21(0.20) | 34.22(8.71) | 2.42(0.35) | 0.20(0.04) | 0.37(0.10) | 30.22(10.84)^a^ |
| CC | 0.40(0.05) | -0.33(0.04) | 1.23(0.12) | 20.52(5.30) | 2.44(0.31) | 0.39(0.07) | 0.24(0.05) | 33.99(7.39)^a^ |
| Rye | 0.35(0.04) | -0.34(0.04) | 1.21(0.11) | 24.69(5.80) | 2.19(0.35) | 0.38(0.06) | 0.26(0.05) | 77.06(14.85)^b^ |

**Figure S2.** Mean difference in predator taxa abundance between cover crop treatments and no-cover conventional treatments in the mid season (pre-emergence and seedling cotton) for 2017 (left) and 2018 (right). Error bars indicate standard error of the mean difference. Predator taxa are ranked by the strongest cover crop effect.

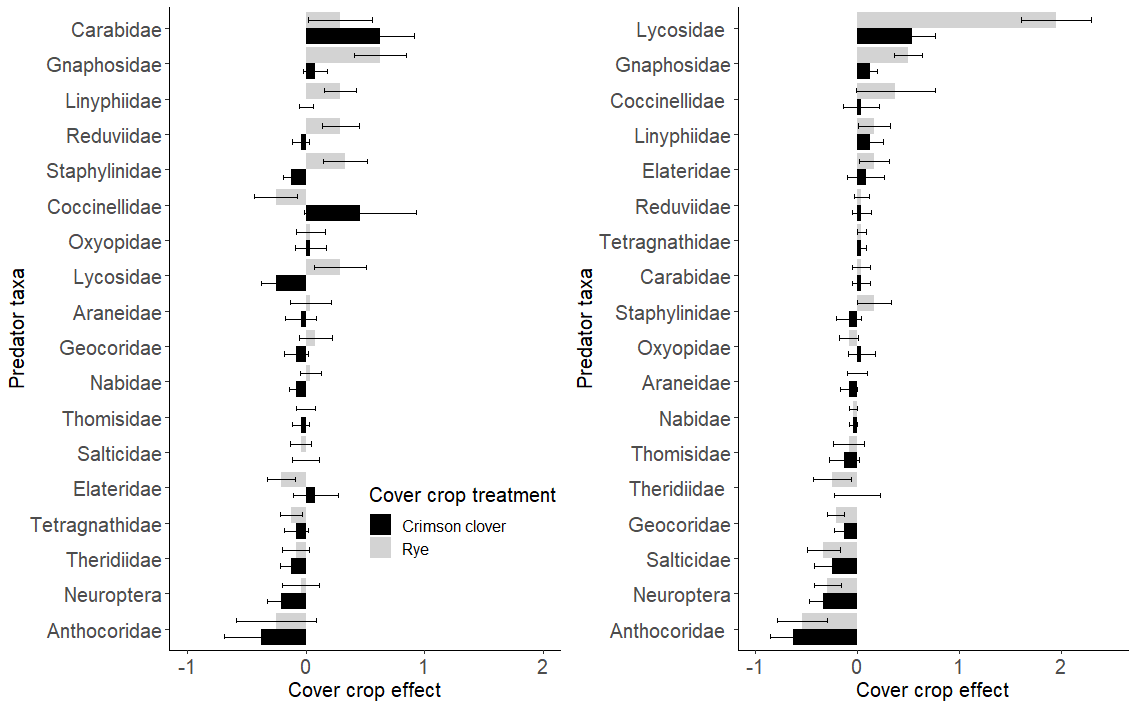

**Figure S3.** Mean difference in predator taxa abundance between cover crop treatments and no-cover conventional treatments in the late season (pre-emergence and seedling cotton) for 2017 (left) and 2018 (right). Error bars indicate standard error of the mean difference. Predator taxa are ranked by the strongest cover crop effect.

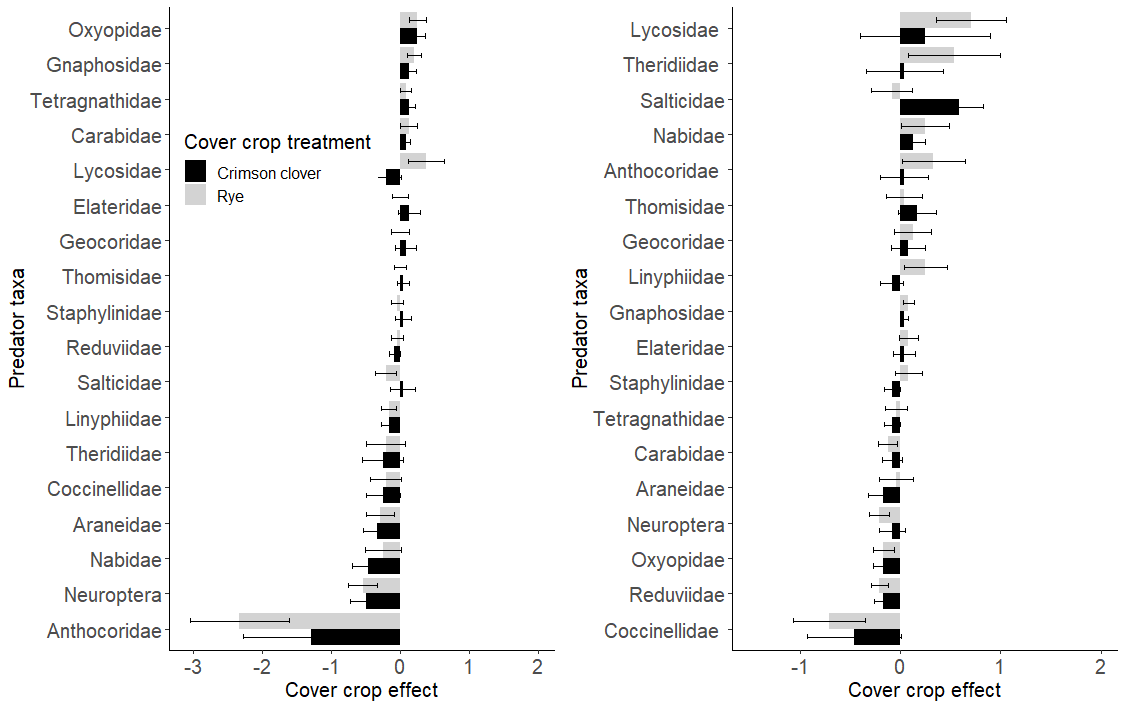

**Figure S4.** Total abundance and composition of predators sampled from each cover crop treatment in the early, middle, and late season for 2017 (A-C) and 2018 (D-F).

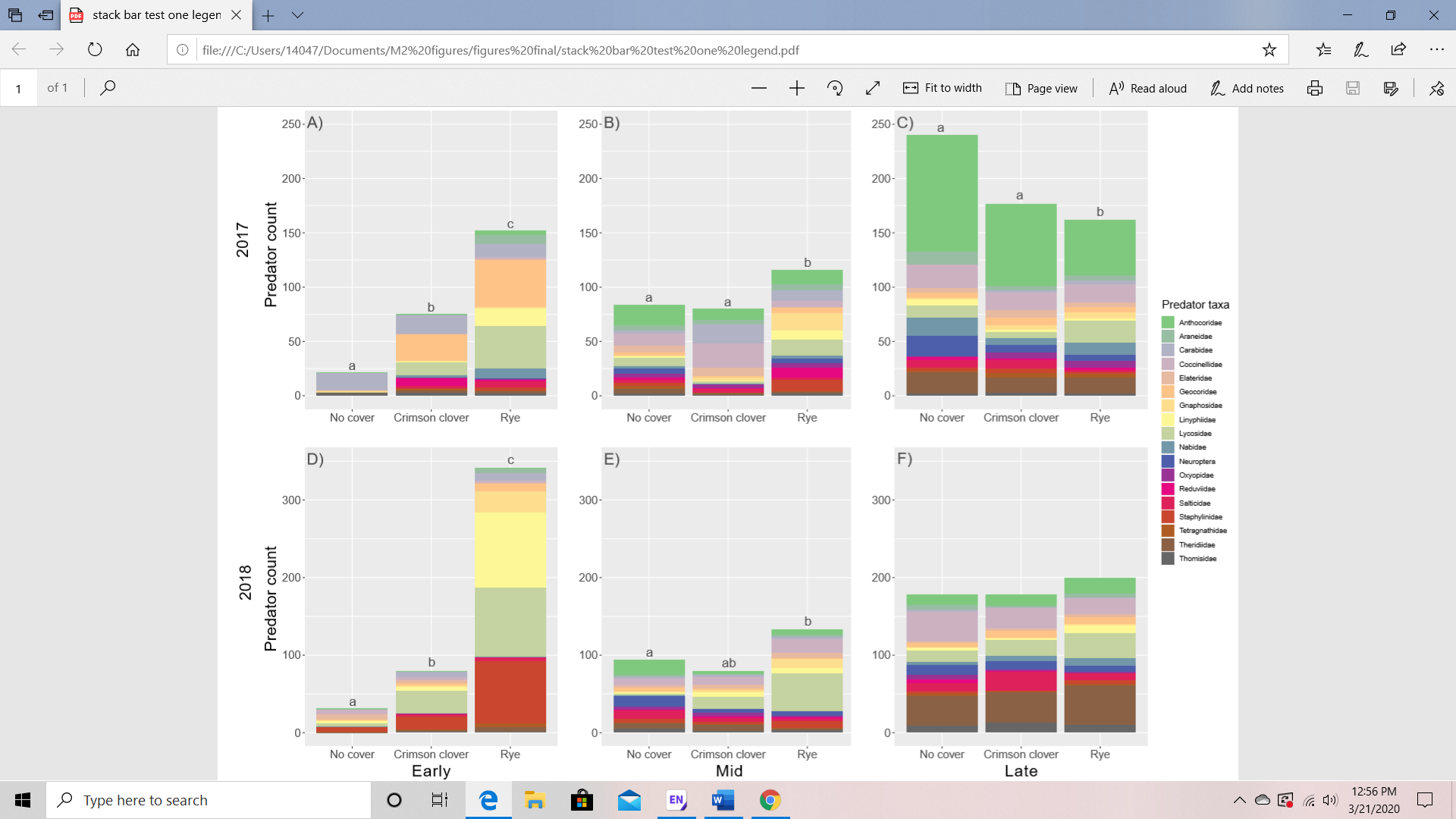
